# The dynamic range of voltage-dependent gap junction signaling is maintained by I_h_-induced membrane potential depolarization

**DOI:** 10.1101/2021.12.16.472972

**Authors:** Wolfgang Stein, Margaret L. DeMaegd, Lena Yolanda Braun, Andrés Vidal-Gadea, Allison L. Harris, Carola Städele

## Abstract

Like their chemical counterparts, electrical synapses show complex dynamics such as rectification and voltage dependence that interact with other electrical processes in neurons. The consequences arising from these interactions for the electrical behavior of the synapse, and the dynamics they create, remain largely unexplored. Using a voltage-dependent electrical synapse between a descending modulatory projection neuron (MCN1) and a motor neuron (LG) in the crustacean stomatogastric ganglion, we find that the influence of the hyperpolarization-activated inward current (I_h_) is critical to the function of the electrical synapse. When we blocked I_h_ with CsCl, the apparent voltage dependence of the electrical synapse shifted by 18.7 mV to more hyperpolarized voltages, placing the dynamic range of the electrical synapse outside of the range of voltages used by the LG motor neuron (−60.2 mV – −44.9 mV). With dual electrode current- and voltage-clamp recordings, we demonstrate that this voltage shift is not due to a change in the properties of the gap junction itself, but is a result of a sustained effect of I_h_ on the presynaptic MCN1 axon terminal membrane potential. I_h_-induced depolarization of the axon terminal membrane potential increased the electrical postsynaptic potentials and currents. With I_h_ present, the axon terminal resting membrane potential depolarized, shifting the dynamic range of the electrical synapse towards the functional range of the motor neuron. We thus demonstrate that the function of an electrical synapse is critically influenced by a voltage-dependent ionic current (I_h_).

**New & Noteworthy:** Electrical synapses and voltage-gated ionic currents are often studied independently from one another, despite mounting evidence that their interactions can alter synaptic behavior. We show that the hyperpolarization-activated inward ionic current shifts the voltage dependence of an electrical synaptic transmission through its depolarizing effect on the membrane potential, enabling it to lie within the functional membrane potential range of a motor neuron. Thus, the electrical synapse’s function critically depends on the voltage-gated ionic current.

## Introduction

Electrical synapses are ubiquitous throughout the nervous system and have been described in many vertebrate and invertebrate systems. Non-rectifying electrical synapses often support neuronal synchrony between neurons of similar functional and anatomical or biochemical profiles through bilateral current flow [1]. In contrast, rectifying synapses are usually found in feedforward neuronal circuits. Their effects are primarily unilateral and current flows preferentially from the presynaptic to the postsynaptic cell. Electrical synaptic transmission shows complex dynamics, including facilitation and depression, and electrical synapse conductance is subject to alterations by a host of influences, including temperature and pH, as well as neuromodulators and hormones [1, 2]. These effects can either be mediated through direct actions on the gap junction proteins, or indirectly through interactions with other electrical processes in neurons, such as voltage-gated ion channels, that affect synaptic current flow and alter the dynamics of the electrical coupling. However, our understanding of such interactions, and their consequences for the electrical behavior of the synapse, is still in its infancy.

Included among the currents influencing electrical synaptic transmission are ionic currents with slow kinetics, such as persistent Na^+^ currents and the hyperpolarization-activated inward current (I_h_). Their slow kinetics make them particularly conducive to changing the properties of electrical synaptic transmission, for example through changes in the extrajunctional membrane resistance or the postsynaptic membrane potential [1, 2]. They thus act without altering the properties of the gap junction proteins themselves, but instead set the condition within which synaptic transmission operates. For instance, the persistent Na^+^ current depolarizes postsynaptic cerebellar Golgi neurons, which causes a switch in the sign of the electrical synapse from predominantly hyperpolarizing to predominantly depolarizing [3, 4]. I_h_ has also been implicated in altering electrical synaptic transmission [5], although examples are less prevalent. In neurons of the snail *Helisoma*, for example, the modulation of electrical synapses between identical neurons is mediated by an I_h_-mediated decrease in extra-junctional membrane resistance [6].

One system that is ideal for studying the effects of I_h_ on electrical synaptic transmission is the crustacean stomatogastric nervous system. Here, several electrical synapses and their effects on the behavior of the motor circuits in the stomatogastric ganglion (STG) have been characterized. In the crab, *Cancer borealis*, the descending modulatory projection neuron 1 (MCN1) innervates the lateral gastric (LG) motor neuron in the STG’s gastric mill central pattern generator via both chemical and electrical synaptic connections [7, 8]. MCN1’s chemical synapse provides slow neuropeptidergic excitation and enables LG to burst and produce rhythmic activity. The electrical synapse contributes to the maintenance of the LG burst [9]. Thus, both chemical and electrical synapses are crucial components required for LG’s behavior. The electrical synapse shows a peculiar voltage dependence. At rest, each MCN1 action potential elicits a small sub-threshold electrical postsynaptic potential (ePSPs) in LG (Fig. 1A, black arrows). However, when LG depolarizes, these ePSPs increase in amplitude. This voltage dependence appears to be a property of the electrical synapse itself.

**Figure 1.**
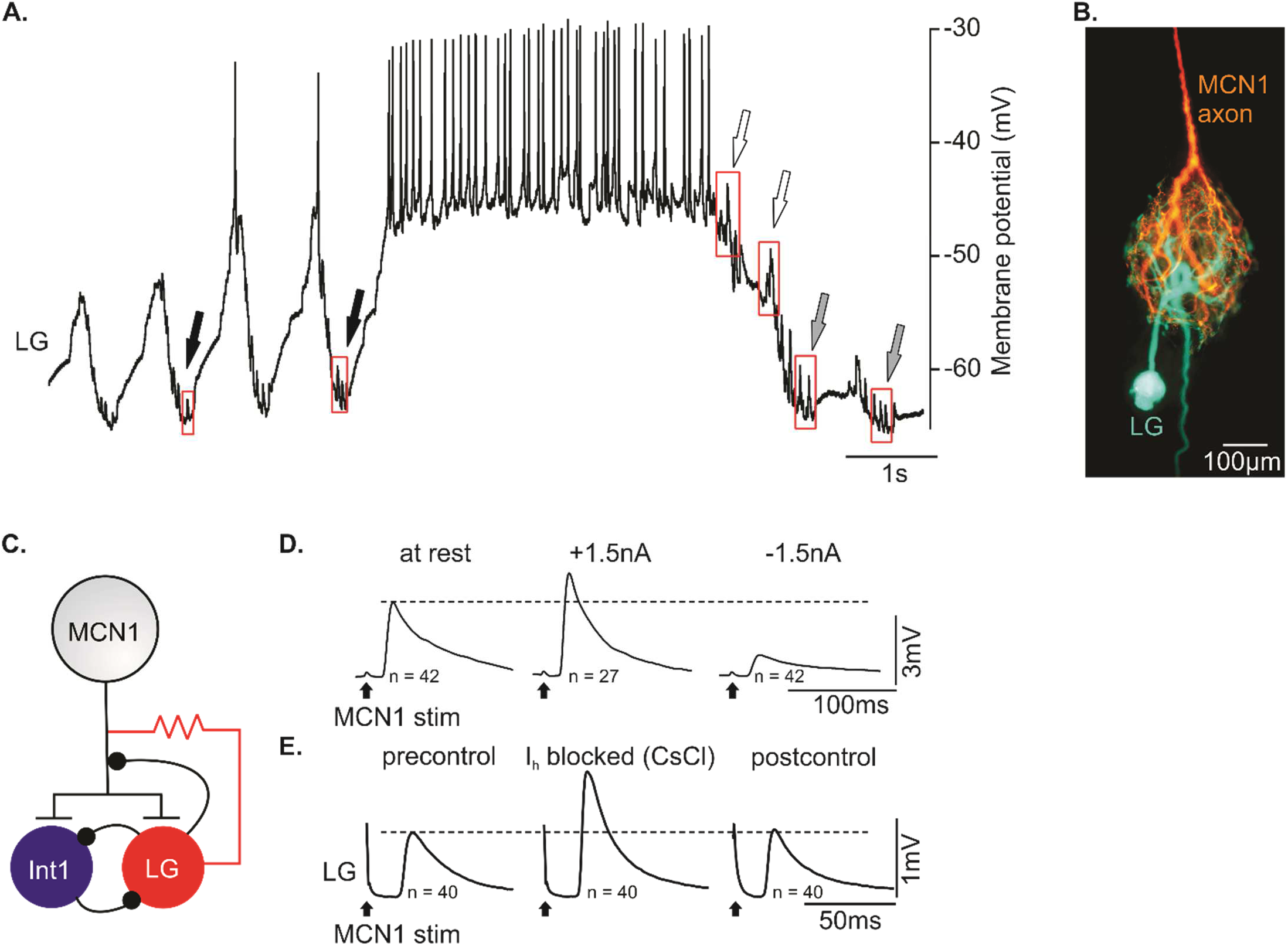
MCN1 elicits voltage-dependent electrical postsynaptic potentials in LG. A. Intracellular recording of LG membrane potential during ongoing gastric mill rhythm. In the LG interburst, ePSPs are small (black arrows). At the end of the LG burst, ePSPs are larger (outlined arrows). ePSP amplitudes slowly diminish after the burst (gray arrows). B. Staining of MCN1 axon terminals (red) and LG cell body and neuropil (green) in the STG. C. Schematic of MCN1 and LG synaptic connectivity. MCN1 chemically excites LG and Int1, which reciprocally inhibit each other. LG provides presynaptic inhibitory feedback to the MCN1 axon terminal. In addition, the MCN1 axon terminal is electrically coupled to LG. D. Time-triggered averages of ePSPs in the absence of a gastric mill rhythm, in low Ca^2+^ saline. Left: at resting membrane potential. Middle: during LG depolarization with 1.5 nA. Right: during LG hyperpolarization with −1.5 nA. Individual ePSPs were elicited through MCN1 axon stimulation with 1 Hz. Averages were calculated by aligning individual sweeps of LG membrane potentials to the MCN1 stimulation. E. ePSP amplitude increases when I_h_ is blocked. Time-triggered averages of MCN1 PSPs in LG. Individual ePSPs were elicited through MCN1 axon stimulation with 1 Hz. Left: in low calcium saline. Middle: with I_h_ blocked by bath application of CsCl (5 mM). Right: washout.

While the basic description of the voltage dependence of the MCN1-LG ePSPs has been available for over 25 years, interactions with (and modulation by) other intrinsic electrical properties of MCN1 and LG are less understood. Here, by quantifying the voltage dependence of the ePSPs and measuring electrical synaptic currents, we show that the ePSPs are not just modulated by the hyperpolarization-activated inward current (I_h_), but that this modulation is critical to their function. Without I_h_ modulation, the voltage dependence of the ePSPs was shifted below physiologically relevant values. As a consequence, ePSPs reached their maximum amplitude at resting membrane potential, but remained invariant during further LG depolarization. Modulation by I_h_ shifted the dynamic range of the ePSP voltage dependence to more positive voltages to match LG’s membrane potential, allowing ePSPs to increase when LG depolarizes. We demonstrate that this voltage shift does not result from a change in the properties of the gap junction, but is due to a sustained depolarization of the MCN1 axon terminal membrane potential that is caused by the presence of I_h_.

## Methods

### Experimental animals

Adult male Jonah crabs (*Cancer borealis*) were obtained from “The Fresh Lobster Company” (Boston, MA, USA) and kept in aerated sea water tanks (~1.025 g / cm^3^ salinity, artificial sea salt, AquaMed, USA). The tanks were kept at a temperature of 10 to 12°C with a 12-hour dark/light cycle. All experiments were performed following the “Principles of animal care” publication No. 86-23, revised in 1985 by the National Institute of Health.

### Dissection

Animals were cold anesthetized on ice for 30 to 40 minutes before dissection. The dissection was carried out following published protocols [10]. In short, the stomach was removed from the animal and the STNS was dissected under a stereomicroscope. During the dissection, the nervous system was constantly kept at a low temperature by rinsing it with 4°C cold saline as appropriate. During experiments a temperature between 10 and 13°C was maintained by superfusing chilled saline.

*Chemicals: Cancer borealis* physiological saline consisted of (in mM, pH 7.4 - 76). KCl 11, NaCl 440, CaCl_2_*2H_2_O 13, MgCl_2_*6H_2_0 26, Trizma base 11.2, Maleic acid 5.1 (all from Sigma Aldrich, USA) dissolved in ultrapure water (18.3MOhm). In some experiments, saline with low calcium concentration (low Ca^2+^- saline) was used to block calcium channels and synaptic release. Low Ca^2+^- saline consisted of (in mM): KCl 11, NaCl 440, CaCl_2_*2H_2_O 0.1, MgCl_2_*6H_2_0 26, Trizma base 11.2, Maleic acid 5.1, MnCl_2_ 12.9. Manganese was added after adjusting the pH to 7.4 – 7.6. To selectively block I_h_, Cesium chloride (CsCl, Sigma-Aldrich, USA) was dissolved in low Ca^2+^- saline (5 mM) before every experiment [11].

### Extracellular recordings

Extracellular recordings followed established protocols [12]. In short, relevant nerves were electrically isolated with petroleum jelly wells. Subsequently, a recording electrode was placed inside the well while a reference electrode was placed in the grounded outside bath. The signal was differentially amplified using an A-M Systems Model 1700 amplifier (Carlsborg, WA, USA) and digitized with a Micro 1401 mkII (Cambridge Electronics Design, Cambridge, UK). Spike 2 (ver. 7, Cambridge Electronics Design, Cambridge, UK) was used to record and analyze the data. To detect the activity of the lateral gastric motor neuron (LG), the lateral gastric nerve (*lgn*) was recorded. It contains the LG axon.

### Intracellular recordings

A dark field condenser (Nikon) was placed in-between LED and Petri dish to facilitate the visual contrast during intracellular recordings. Glass microelectrodes (borosilicate glass capillaries GB100TF-8P, Science Products GmbH, Hofheim, Germany) were pulled with a microelectrode puller (Model P-97, Sutter Instrument, Novato, USA) and filled with a 0.6 M K_2_SO_4_ solution. Recordings with obvious changes in membrane voltage or cell input resistance over the course of the experiment were discarded. Electrode resistance ranged between 15-25 MΩ. To impale cells, either a manual micro-manipulator (Leitz, Wetzlar, Germany) or several electrical manipulators (Sutter, Novato, USA) were used to position the electrodes above the STG. LG was identified by comparing its intracellularly recorded action potentials to those on the extracellular *lgn* recording. MCN1 was identified by responding to stimulation of its axons in the inferior esophageal nerve (*ion*; see stimulation below). Signals were amplified through either Axoclamp-2B, Axoclamp 900B (Molecular Devises, Sunnyvalle, CA, USA) or NPI BA-1s (NPI Electronic GmbH, Tamm, Germany) amplifiers. For two-electrode current or voltage clamp recordings, two electrodes were simultaneously inserted into LG or the MCN1 axon terminal.

### Neuronal stainings

To visualize the MCN1 axon terminals and the LG neuropil (Fig. 1B) we iontophoretically injected neurons with a fluorescent dye. Alexa Fluor 568 (MCN1 terminal) and Alexa Fluor 488 (LG) potassium salts (Thermo Fisher Scientific, Waltham, MA, USA) were diluted in 200 mM KCl and loaded in the electrode tip. After dye loading, electrodes were backfilled with 200 mM KCl. Negative current pulses of −10nA were applied for at least 10 minutes (2 seconds on, 0.5 seconds off) to load cells with dye.

### Extracellular nerve stimulation

Petroleum jelly wells were placed around each *ion* to stimulate the axon of MCN1, following established protocols [13]. In some experiments the *ions* were cut close to the commissural ganglia that house the somata of the two MCN1 neurons. To activate MCN1, the remaining section of one *ion* still attached to the STG was stimulated. 1 Hz stimulation with a duration of 1 ms were applied elicit individual action potentials [14]. The long inter-pulse interval prevented a temporal summation of chemical PSPs in LG.

### Model

We used a model consisting of two electrically coupled point neurons – one presynaptic and one postsynaptic. For each neuron, the membrane potential *Vm* is found from

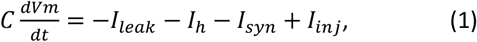

where *C* is the capacitance, *I_L_* is the leak current, *I_h_* is the hyperpolarization-activated inward current, *I_syn_* is the synaptic current, and *I_inj_* is the injected current.

The leak current is given by

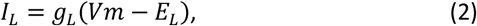

where *g_L_* is the leak conductance and *E_L_* is the leak equilibrium potential.

The hyperpolarization-activated inward current is given by

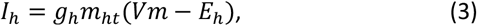

where *g_h_* is the conductance, *m_ht_* is the time- and voltage-dependent activation function, and *E_h_* is the equilibrium potential.

The time-dependence of the activation function is found by solving the differential equation

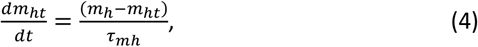

where the voltage dependence of the activation function is given by

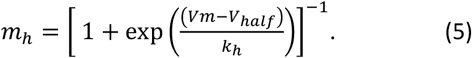

The activation function time constant is *τ_mh_* and the midpoint and growth rate of the logistic function are *V_half_* and 1/*k_h_* respectively.

The synaptic current is given by

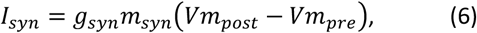

where *g_syn_* is the conductance, *m_syn_* is the voltage-dependent activation function, and *Vm_prh,post_* are the pre- and postsynaptic membrane potentials, respectively.

The synaptic activation function is given by

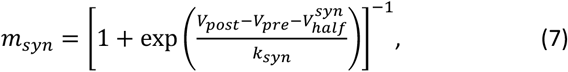

where 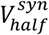 and 1/*k_syn_* are the midpoint and growth rate respectively.

The injected current is given by a square pulse of duration *t_pulse_* and height *I_pulse_* for the presynaptic neuron. For the postsynaptic neuron, the injected current is a constant that is varied between −2 nA and +38 nA in the simulation to achieve different membrane potentials.

All model parameters are listed in Table 1

**Table 1.**
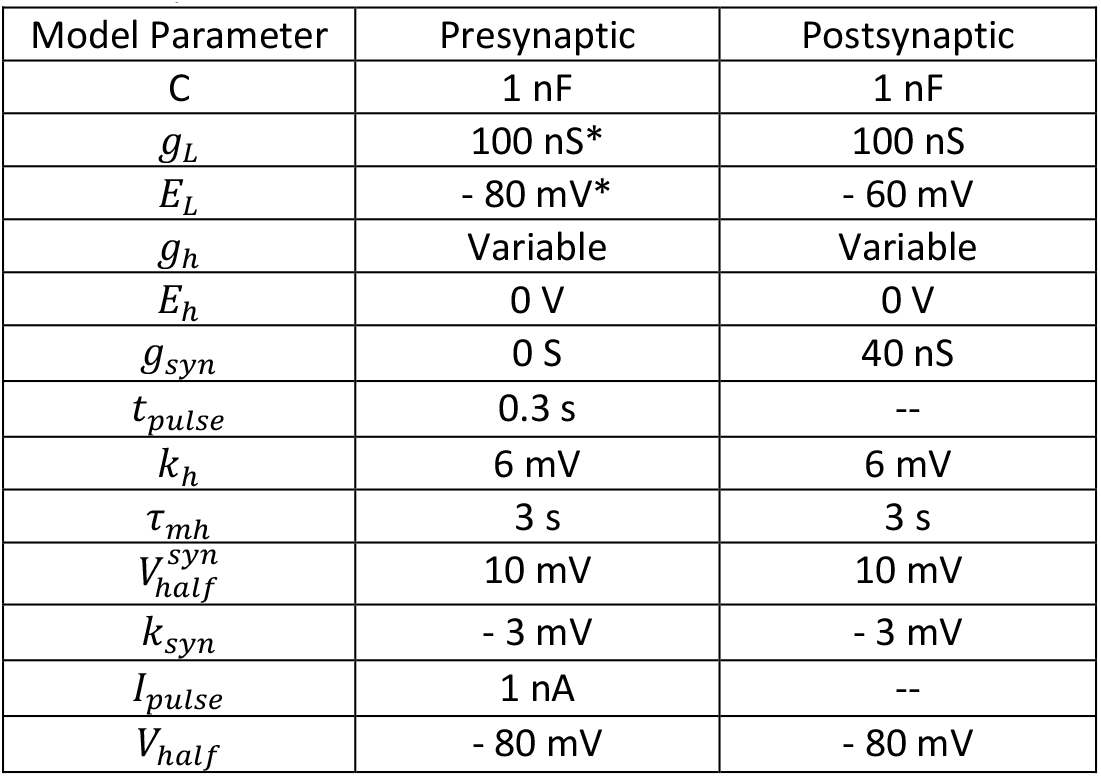
Model parameters. *default parameters used unless otherwise stated in the text.

The simulation was performed with a novel code in Wolfram Mathematica v12.1 (RRID:SCR_014448) using the NDSolve command for the differential equations. The model is available at ModelDB (accession number 267286). The parameters were selected to represent physiologically meaningful values. No specific tuning was necessary, i.e., parameters were not tuned to represent the biological neurons (MCN1 and LG). Neuronal behavior was observed for 30 seconds with the presynaptic pulse initiated at *t* = 10 s. The postsynaptic pulse was detected by using the NMaximize function in Mathematica for 10.2 ≤ *t* ≤ 11 with a working precision of 10. Initial conditions for solving the differential equations are listed in Table 2.

**Table 2.** Initial conditions used to solve the differential equations in Eqs. (1) and (4).

| Function | Initial Condition |
| --- | --- |
| $Vm_{pre}$ | - 70 mV |
| $Vm_{post}$ | - 60 mV |
| $m_{ht_{pre}}$ | 0.5 |
| $m_{ht_{post}}$ | 0.5 |

### Analyses and visualizations

Data were recorded and analyzed with Spike 2 script language and built-in functions. Measurements were transferred for spreadsheet analysis to Excel (Version 365 for Windows, Microsoft). Final figures were generated using Corel-Draw (Version X7 for Windows, Corel Corporation, Ottawa, ON, Canada). Statistical tests were carried out with SigmaPlot (Version 11 for Windows, Systat Software GmbH, Erkrath, Germany, RRID:SCR_003210). Sigmoidal fits were generated using the SigmaPlot Nonlinear Regression - Dynamic Fitting function, using the equation f= a/(1+exp(-(x-x0)/b)) and 200 iterations. Here, x0 represents the midpoint of the sigmoidal fit that was used to compare between low Ca^2+^ saline and low Ca^2+^ saline with added CsCl (5mM). Only data with significant regression (p<0.05) were used. Individual results are given in the text or the figure legends. “N” denotes the number of animals tested whereas “n” indicates the number of repetitions used in an experiment.

## Results

### The electrical synapse between MCN1 and LG is voltage-dependent

MCN1 is part of a group of descending projection neurons located in the commissural ganglia (CoGs) [15]. There is one MCN1 neuron in each of the two CoGs and both project several centimeter-long axons to the STG via the inferior esophageal (*ion*s) and stomatogastric nerves (*stn*). After entering the STG neuropil (Fig. 1B), the MCN1 axons innervate pyloric and gastric mill neurons. While the concurrent activity of both MCN1 copies appears important to elicit robust gastric mill activity [16], the two MCN1 axons have identical postsynaptic targets. Each MCN1 axon terminal interacts via several direct synaptic connections with the core gastric mill CPG neuron LG (Fig. 1C). MCN1 excites LG chemically via *Cancer borealis* tachykinin-related peptide Ia (CabTRP Ia) [14]. The slow LG membrane potential depolarization resulting from this chemical excitation eventually activates spike generation in LG, allowing it to escape the inhibition from its antagonist, Interneuron 1 (Int1). LG also has a chemical feedback synapse onto the MCN1 axon terminal, which over time weakens the chemical excitatory input LG obtains from MCN1 [9].

In addition, the MCN1 axon terminal and LG are electrically coupled. MCN1 action potentials elicit subthreshold ePSPs that are readily observable in LG (Supplemental Figure S1A (https://doi.org/10.6084/m9.figshare.17303918.v1)). Fig. 1A highlights several ePSPs during an ongoing gastric mill rhythm (red boxes, arrows). While the interactions between the chemical synapses allow LG to create its burst, the ePSPs contribute to burst maintenance by prolonging burst duration [9, 17]. The electrical synapse between the MCN1 axon terminal and LG appears unaffected by the presynaptic inhibition LG exerts on the axon, but the elicited ePSPs show an obvious voltage dependence: they grow in amplitude when LG depolarizes. Fig. 1A shows this during an ongoing gastric mill rhythm. When LG depolarizes and starts to burst, ePSPs increase in amplitude, prolonging the burst with action potentials riding on top of the ePSPs. This can be seen best by comparing ePSPs before the LG burst (black arrows) to those during and immediately after the burst (outlined arrows). When the LG burst ends due to a reduction of MCN1 neuropeptide release [18], the LG membrane potential hyperpolarizes and the amplitude of the ePSPs decreases again (Fig. 1A, gray arrows after the burst; [9]). The voltage dependence of the ePSPs can also be observed without a gastric mill rhythm, and when chemical release from MCN1 and chemical synapses from other STG neurons (such as Interneuron 1; Fig. 1C) are blocked. Fig. 1D shows averaged ePSPs obtained from an intracellular recording of LG’s membrane potential during continuous 1 Hz stimulation of MCN1’s axon (see Methods). No gastric mill rhythm was elicited, and chemical synaptic transmission was blocked using low Ca^2+^ saline. When LG was depolarized by current injection, ePSP amplitude increased. In contrast, when LG was hyperpolarized by a similar current injection, ePSP amplitude diminished below control levels. The voltage-dependent changes in ePSP amplitude are thought to be caused by a voltage-dependence of the electrical synapse itself [9]. We noted, however, that the ePSP amplitude and voltage dependence appeared to be altered by I_h_. Fig. 1E shows averaged ePSPs obtained from an intracellular recording of LG’s membrane potential during continuous 1 Hz stimulation of MCN1’s axon (see Methods). For all conditions shown, chemical synaptic transmission was blocked with low Ca^2+^ saline [19]. When we additionally blocked I_h_ by adding extracellular CsCl (5mM) [20], ePSP amplitudes increased. This suggested that I_h_ intrinsic to either MCN1’s axon terminal or to LG influences the electrical synaptic transmission between them.

### The hyperpolarization-activated inward current (I_h_) shifts the voltage dependence of the MCN1-LG electrical postsynaptic potentials

To investigate I_h_‘s influence, we compared the voltage dependence of the MCN1-LG ePSPs before and after blocking I_h_. Figure 2A (left) shows original ePSP recordings at different LG membrane potentials. Individual MCN1 action potentials were elicited through 1 Hz stimulation of one *ion*, and chemical synapses were blocked with low calcium saline. PSPs were recorded with an intracellular electrode in LG’s soma and LG membrane potential was altered with current injection through a second electrode. As expected, depolarizing LG increased ePSP amplitude, and hyperpolarizing LG decreased ePSP amplitude. We noted, however, that the change in ePSP amplitude was not instantaneous. Instead, it took several hundred ms (477 ± 281 ms, N=7) after a voltage step for the amplitudes to reach their new equilibrium (Supplemental Figure S1B (https://doi.org/10.6084/m9.figshare.17303918.v1)).

**Figure 2.**
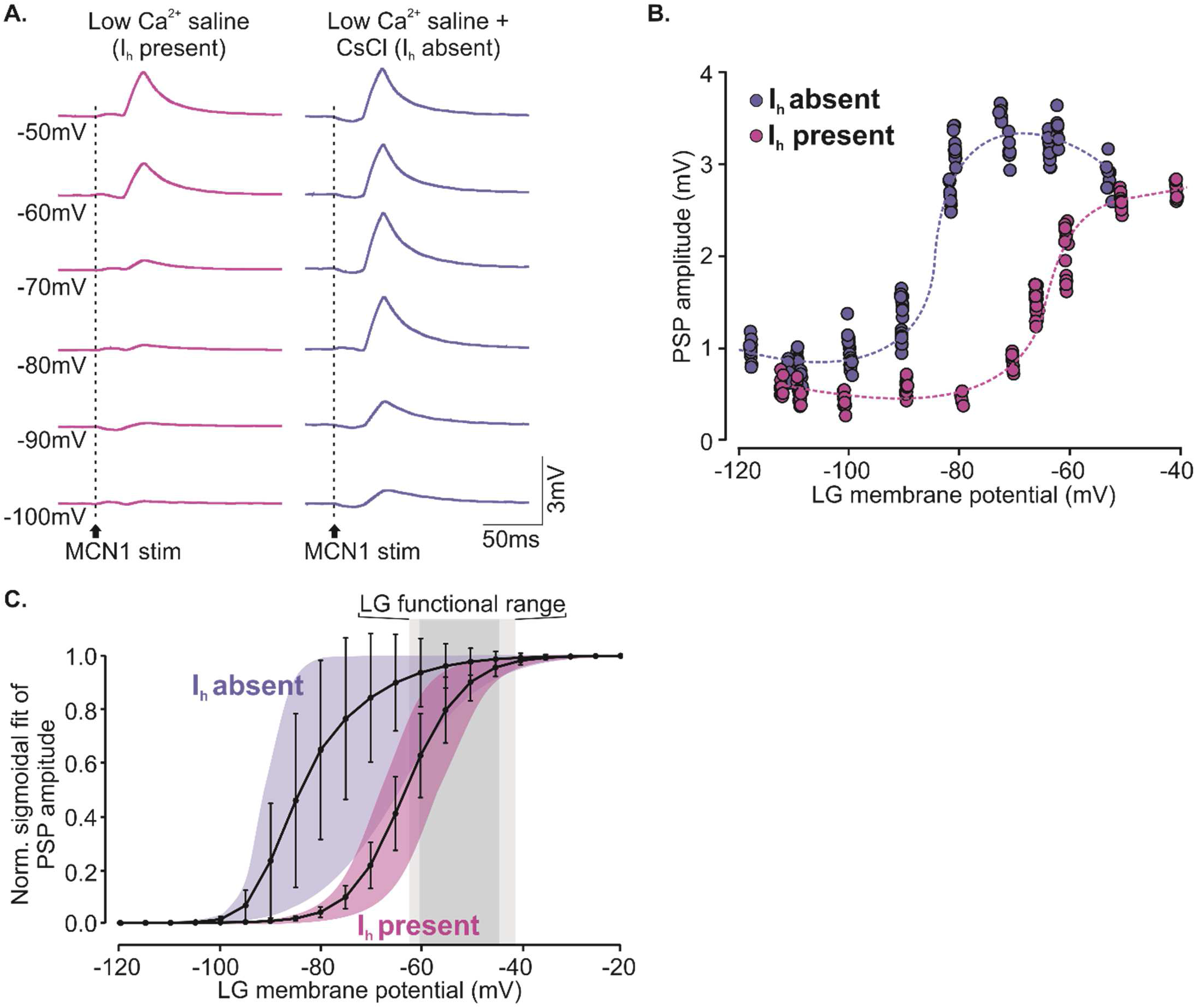
Blocking I_h_ shifts the voltage dependence of the MCN1 ePSPS to more hyperpolarized voltages. A. Original traces of MCN1 ePSPs at various LG membrane potentials (−50 to −100 mV). In CsCl (right) ePSPs were larger, and only diminished at more hyperpolarized LG membrane potentials. Individual ePSPs were elicited through MCN1 axon stimulation with 1 Hz. B. Quantification of ePSP amplitudes of the experiment shown in (A). Multiple ePSPs were elicited at each membrane potential. Magenta: in low calcium saline. Purple: with I_h_ blocked by CsCl. C. Quantification of all experiments. The ePSP voltage dependence of each experiment was fitted with a sigmoidal curve. Mean ± SD for all experiments are shown. Colored areas indicate the range of all observed data. Magenta: in low calcium saline. Purple: with I_h_ blocked by CsCl. The gray box indicates the functional range of LG’s membrane potential. Light gray indicates SD.

Overall, we obtained a mostly sigmoidal relationship between membrane potential and ePSP amplitude (Fig. 2B, magenta), with ePSP amplitudes flattening out towards their smallest values around −70mV, and towards their maximum at around −40mV. The midpoint of the sigmoidal curve was −63.5 mV. Similar results were obtained in all tested animals. Fig. 2C (magenta) shows a summary of all experiments (N= 8 animals), with the averaged sigmoidal of all experiments, and their standard deviation. On average, the midpoint of the ePSP amplitude change was −62.7±3.3 mV (for calculation of the sigmoidal fits see Methods). When we blocked I_h_, ePSP amplitude generally increased, but most dramatically at voltages near the LG resting membrane potential (−75mV in this particular experiment). Fig. 2A (right) shows the resulting ePSP amplitudes for one experiment, and the resulting sigmoidal is shown in Fig. 2B (purple). In this experiment, the midpoint of the sigmoidal had shifted to – 86.7 mV and was thus more hyperpolarized than with I_h_ present. We found similar results in all experiments (Fig. 2C, purple). On average, the midpoint of ePSP amplitude change was now −81.4±9.0 mV (N=7, Mann-Whitney U-test, P=0.002, significantly different from no CsCl control), and amplitudes already approached their maximum around −70mV. Thus, the dynamic range of the ePSP voltage dependence appeared to be no longer within the functional range of the LG membrane potential. Indeed, when we measured the range of subthreshold membrane voltages LG exhibits during a gastric mill rhythm, we found that, on average, the resting membrane potential was −60.2 ± 2.3 mV, the most hyperpolarized membrane potential during an ongoing rhythm was −59.3 ± 2.0 mV, and the spike threshold was −44.9 ± 3.2 mV (N=14). We gathered these data in a different set of experiments, in regular saline without blockers to allow the gastric mill rhythm to occur. Comparing LG’s subthreshold voltage range to the voltage dependence of the electrical synapse (Fig. 2C) revealed that 33% of the ePSP amplitude change occurred in that voltage range when I_h_ was present. In contrast, with I_h_ blocked, the amplitude changes occurred at more hyperpolarized membrane potentials, and less than 9% occurred within LG’s functional voltage range. This suggests that I_h_ plays a critical role in keeping the dynamics of this synapse in the adequate voltage range.

### I_h_ can be detected in both LG and the MCN1 axon terminal

While many STG neurons possess I_h_, it has not been studied in detail in either LG or MCN1. As a first step, we used two-electrode current clamp to test if I_h_ was present. Fig. 3A shows an example recording in which LG was hyperpolarized through current injection for several seconds and then released from hyperpolarization. The sag potential (arrows in Fig. 3A) typical for I_h_ is clearly visible and slowly depolarized LG during the current step. After release from hyperpolarization, LG’s membrane potential overshot the resting membrane potential in a postinhibitory rebound, but only by a few mV. The recording shows small spikelets (2 mV and smaller) that ride on top of the rebound (asterisk in Fig. 3A). These spikelets were not LG action potentials. Instead, these were ePSPs originating from MCN1, which apparently went through its own rebound, elicited by the strong hyperpolarization of LG. In more than 10 LGs in which we caried out similar current injections, only one showed a rebound that elicited LG action potentials. This suggested that I_h_ was either weak or absent in LG, but present in the MCN1 terminal, and our current injection reached MCN1 through the gap junction. Fig. 3B shows a step protocol where LG was hyperpolarized in multiple steps. I_h_ was activated, but again only at very hyperpolarized membrane potentials (<−90 mV), using rather large injected currents (>-20 nA). Fig. 3C shows the same step protocol after CsCl application. Here, no sag potential or rebound was present, confirming that indeed I_h_ was recorded. Figures 3D & 3E show the same scenario in the two-electrode voltage clamp with similar results. In control (low calcium saline), I_h_ was detectable but absent in CsCl. This was consistent across experiments (N=7 for current clamp, N=11 for voltage clamp). We noted, however, that in all experiments, I_h_ was only activated at rather negative membrane potentials (see discussion).

**Figure 3.**
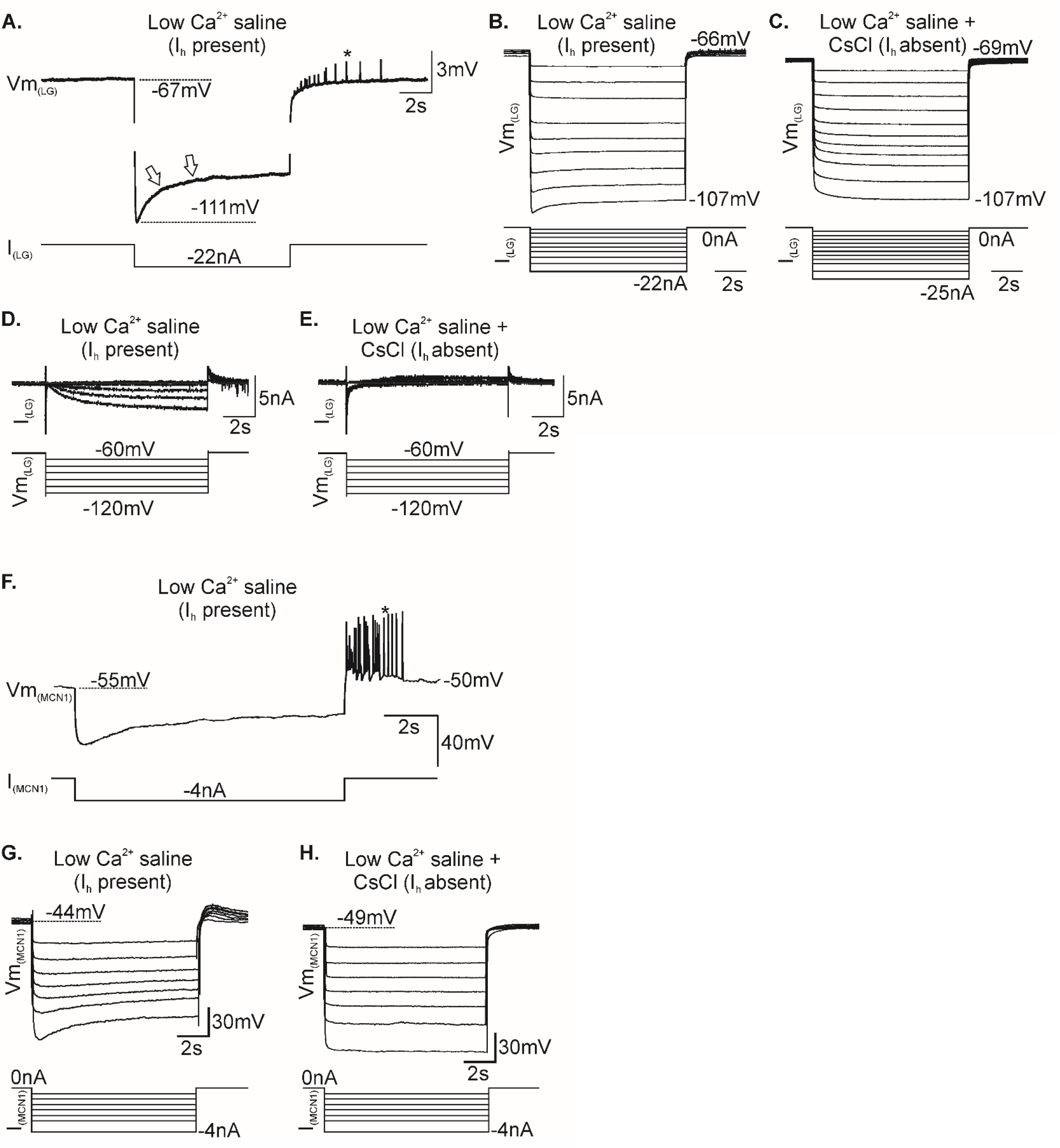
The hyperpolarization-activated inward current can be observed in LG and in the MCN1 axon terminal. A. Two-electrode current clamp recording of LG. When LG was hyperpolarized by current injection, the membrane potential showed a slow sag during which the voltage depolarized. After the end of the hyperpolarization, there was a small rebound as the membrane potential surpassed the resting membrane potential. No LG spikes were generated. *ePSPs elicited by MCN1 action potentials during the rebound. Arrows: sag potential. B. Step protocol in two-electrode current clamp in low calcium saline. A voltage sag was visible below −90 mV. C. Step protocol in two-electrode current clamp in low calcium saline, but with I_h_ blocked by CsCl. No voltage sag was detectable. D. Two-electrode voltage clamp recording with step protocol hyperpolarizing the LG membrane potential, in low calcium saline. A small I_h_ was visible. E. Same step protocol in CsCl. I_h_ was absent. F. Two-electrode current clamp recording of the MCN1 axon terminal. When the terminal was hyperpolarized by current injection, the membrane potential showed a sag during which the voltage depolarized. After the end of the hyperpolarization, MCN1 rebounded and produced several action potentials (*). G. Current step protocol to hyperpolarize the MCN1 axon terminal in low calcium saline. A clear sag of the membrane potential was visible. H. Same protocol, but with I_h_ blocked by CsCl. The voltage sag and the rebound after the end of the voltage steps were absent.

To determine if I_h_ was present presynaptically, we recorded intracellularly from the MCN1 axon terminal. Some of these recordings were done with a single electrode due to the difficulty to find the MCN1 axon and impale it with two electrodes. Nevertheless, we found evidence that I_h_ is present in the MCN1 axon terminal. Fig. 3F shows an example of a two-electrode recording where we hyperpolarized the MCN1 axon terminal for several seconds. The membrane potential slowly depolarized during the current step. After the end of the current injection, MCN1 rebounded and fired several action potentials in a response that was much stronger than in our LG recordings (compare Fig. 3A). Notably, these action potentials were generated in the axon terminal and were not reflections of possible postsynaptic action potentials in LG since no action potentials occurred on the extracellular LG recording (not shown). Fig. 3G shows a step protocol where MCN1 was hyperpolarized in a single electrode configuration. As such, currents, rather than membrane potentials, are given for the step protocol. Even small current injections (4nA and smaller) were sufficient to elicit clear sag potentials. This was not unexpected since current was injected into a small-spaced axon, as opposed to an entire neuron in the case of LG. In the presence of CsCl, the MCN1 sag potentials, as well as the rebound of the membrane potential after release of the current injection, were absent (Fig. 3H), demonstrating that they were caused by I_h_. This was the case in all preparations (N=6).

### I_h_ shifts the observed voltage dependence of an electrical synapse by depolarizing the presynaptic membrane voltage

To address the question how I_h_ may affect the voltage dependence of the MCN1-LG ePSPs, we built a computational model of two neurons connected via a voltage-dependent electrical synapse. In this simplistic model (see Methods for details and default parameters used), each neuron consisted of only one compartment. The electrical synapse only passed current in one direction (from the presynaptic to the postsynaptic neuron). The voltage dependence was described by equation 7 (see Methods), which depends not just on the presynaptic membrane potential, but also on the postsynaptic membrane potential. This sigmoidal activation function resulted in a sigmoidal amplitude increase of the postsynaptic potentials when the postsynaptic membrane potential was depolarized (Fig. 4A, left: model output, Figs. 4B & 4C, red traces of sigmoidal response plot). Both neurons possessed passive membrane properties, but no active sodium or potassium currents (and therefore no action potentials) as those are not necessary to measure synaptic events. A single current pulse into the presynaptic neuron was used to elicit rapid changes in the membrane potential, simulating an action potential. We additionally implemented I_h_, either in the presynaptic neuron or the postsynaptic neuron, or both. We varied the amount of I_h_ via its maximum conductance (g_h_) from 20 to 80 nS. Results were compared to a control condition without I_h_ (g_h_= 0 nS).

**Figure 4.**
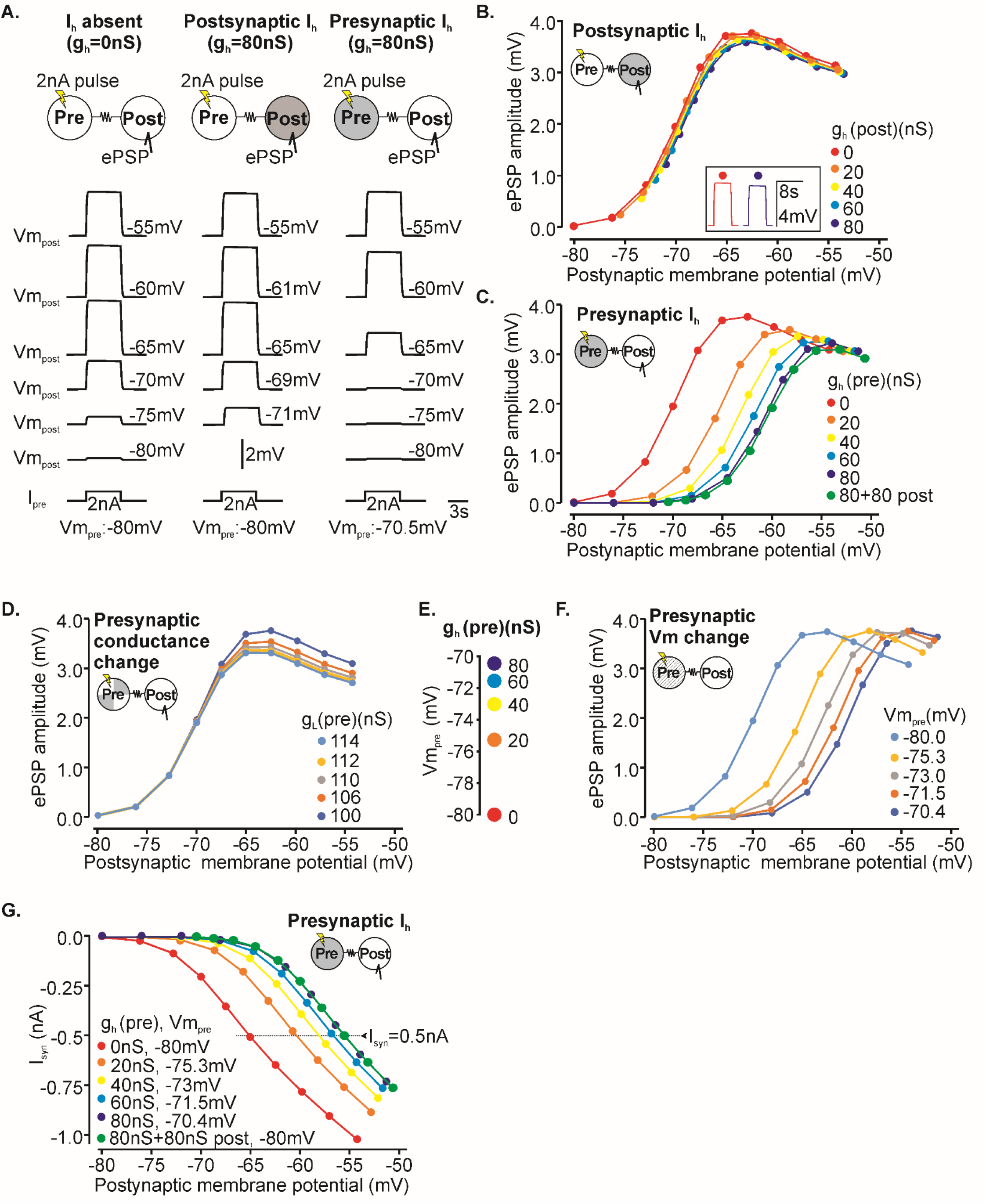
Modelling predicts that I_h_-induced changes to the presynaptic membrane potential shift the voltage dependence of the ePSPs. A. Electrical postsynaptic potentials at various membrane potentials of the postsynaptic neuron. Without I_h_ (left), the largest increase of ePSP amplitude occurred between −75 and −70 mV. We obtained very similar results with I_h_ in the postsynaptic neuron (middle). Note, the most hyperpolarized postsynaptic membrane potential was −71 mV because I_h_ depolarizes the membrane potential. With I_h_ in the presynaptic neuron (right), the largest increase occurred between −65 and −60 mV. Top: schematic of stimulation and recording conditions. Vm = membrane potential. Bottom: the resting membrane potential of the presynaptic neuron is given. B. Quantification of ePSP amplitudes with changing postsynaptic membrane potential. The colors indicate different levels of I_h_ in the postsynaptic cell (g_max_ = 0 – 80 nS). The voltage dependence of the ePSP was not affected by I_h_ in the postsynaptic cell, but the maximum amplitudes were diminished slightly. Inset: largest ePSPs elicited without I_h_ and with maximum I_h_ (g_max_ = 80 nS). C. Quantification of ePSP amplitudes with changing postsynaptic membrane potential. The colors indicate different levels of I_h_ in the presynaptic cell (g_max_ = 0 – 80 nS). The voltage dependence of the ePSPs shifted to more depolarized voltages with more I_h_ and the maximum amplitudes diminished. Adding maximum I_h_ (g_max_ = 80 nS) to the postsynaptic cell (green) did not further change the voltage dependence. D. Quantification of ePSP amplitudes with changing postsynaptic membrane potential. The colors indicate different levels of leak current in the presynaptic cell (g_leak_ = 100 – 114 nS). The voltage dependence of the ePSPs was unaffected, but the maximum amplitudes diminished with more leak. E. Resting membrane potential changes with different levels of I_h_ in the presynaptic cell (g_max_ = 0 – 80 nS). More I_h_ depolarized the presynaptic membrane potential. F. Quantification of ePSP amplitudes with changing postsynaptic membrane potential. The colors indicate different resting membrane potentials (−80 to −70.4 mV). The voltage dependence of the ePSPs shifted to more depolarized voltages with more depolarized membrane potentials in the presynaptic cell. No changes to the maximum amplitudes were observed. G. Quantification of synaptic current (I_syn_) with changing postsynaptic membrane potential. The colors indicate different levels of I_h_ in the presynaptic cell (g_max_ = 0 – 80 nS) and the associated presynaptic membrane potentials (Vm_pre_). With more I_h_, the observed activation of the synaptic current shifted to more depolarized voltages. Adding maximum I_h_ (g_max_ = 80 nS) to the postsynaptic cell (green) did not further change the synaptic current. The dashed line indicates −0.5 nA of synaptic current. Synaptic current is plotted as an inward current with negative sign.

Without I_h_, ePSP amplitudes started to visibly increase at −75 mV and approached a maximum near −65 mV in a sigmoidal curve and a midpoint voltage of −70 mV. Amplitudes decreased slightly at more depolarized membrane voltages. Fig. 4A (left) shows the model output at selected postsynaptic membrane potentials and the resulting sigmoidal change of the ePSP amplitudes is shown in Fig. 4B (red trace). With I_h_ in the postsynaptic neuron, the obtained voltage dependence of the ePSPs remained unchanged, regardless of I_h_ amount (Fig. 4A, middle: model output at selected membrane potentials; Fig. 4B, various colors: sigmoidal changes in ePSP amplitudes), but there was a small overall decrease in ePSP amplitude. The inset in Fig. 4B shows the highest amplitudes achieved without I_h_ vs. the strongest I_h_ (g_h_= 80 nS). In contrast, when I_h_ was implemented in the presynaptic neuron, but not the postsynaptic one (Fig. 4A, right: model output), ePSP amplitudes decreased notably with more I_h_, and the ePSP voltage dependence started to shift toward more depolarized values (Fig. 4C, various colors). At a g_h_ of 80 nS, the midpoint had reached −62 mV, without dramatic changes to the sigmoidal shape of the curve. These results matched our experimental data where in the presence of I_h_, ePSP amplitudes were smaller than when I_h_ was blocked and the midpoint of the curve was more depolarized with I_h_ (compare to Fig. 2). We obtained no further changes to the behavior of the ePSPs when we additionally added I_h_ to the postsynaptic neuron so that it was present in both neurons (Fig. 4C, green), except a small reduction in the maximum amplitude. This further suggested that I_h_ in the presynaptic, but not the postsynaptic neuron, was responsible for the shift of the ePSP amplitude curve.

How did I_h_ exert its effects on the ePSPs? The model shows that there are two main effects I_h_ has on the presynaptic membrane. One is an overall increase in membrane conductance. I_h_ may thus act to shunt the presynaptic voltage change. Second, I_h_ is an inward current (reversal potential of 0 mV) and hence depolarizes the presynaptic membrane potential. To separate the impact of these two effects on the ePSP amplitudes, we altered them independently of one another in the presynaptic model neuron. First, we kept the membrane potential constant but increased overall conductance. We chose the resting potential without I_h_ as our reference potential (−80 mV). Membrane conductance was increased by adding conductance to the leak current, which has a reversal potential of the reference potential. To estimate the increase in conductance for the leak, we calculated the steady state difference in membrane conductance for each level of I_h_ we had used in the original model (Supplemental Table T1 (https://doi.org/10.6084/m9.figshare.17303936.v1)). The original leak conductance (g_L_) was 100 nS. The new values were: g_h_=20 nS, g_L_ =106.25 nS; g_h_=40 nS, g_L_ =109.51 nS; g_h_=60 nS, g_L_ =111.77 nS; g_h_=80 nS, g_L_ =113.51 nS. Adding the additional conductance, but not I_h_ itself, and keeping the presynaptic membrane potential at −80 mV, we found that ePSP amplitudes generally diminished when more conductance was added (Fig. 4D). However, there was no shift in their voltage dependence.

In contrast, when we kept the membrane conductance the same, but altered the membrane potential of the presynaptic model neuron, we found that the voltage dependence shifted in the predicted direction. Here, instead of activating I_h_, we altered the resting membrane potential of the presynaptic neuron. This was achieved through changing the reversal potential of the general leak current in this neuron. This did not add additional conductance, but readily changed membrane potential. We compared models with resting membrane potentials corresponding to the same I_h_ conductance used as in previous models (g_h_ of 0, 20, 40, 60, and 80 nS). We obtained membrane potentials of −80.0, −75.3, −73.0, −71.5, and −70.4 mV, respectively (Fig. 4E; Supplemental Table T1 (https://doi.org/10.6084/m9.figshare.17303936.v1)). We found that at the more depolarized membrane potential of the presynaptic cell, the voltage dependence of the ePSP amplitudes shifted to more depolarized membrane voltages as well (Fig. 4F). This occurred without an overall change to the maximum ePSP amplitudes.

Thus, the model predicts that the shift in the apparent voltage dependence of the ePSPs is due to a shift of the presynaptic membrane potential and can occur without changes to the gap junction voltage dependence itself. As I_h_ increases, and the presynaptic voltage depolarizes, the difference between the pre- and postsynaptic membrane potentials becomes smaller. This reduces the driving force for currents across the synapse as well as synaptic activation since both depend on the difference between pre- and postsynaptic membrane potentials (equations 6 and 7). To test this prediction, we measured the synaptic current at different levels of presynaptic I_h_ conductance (Fig. 4G) and plotted it over the same range of postsynaptic membrane potentials as in Fig. 4C. As expected from equation 6, the synaptic current at each level of I_h_ (various colors) initially increased on a sigmoidal trajectory and then continued linearly as the synapse was fully activated, but the driving force continued to increase linearly. With more I_h_ and thus more depolarized presynaptic membrane potentials, the activation of the synaptic current shifted toward more depolarized postsynaptic values. For example, without I_h_ (g_h_ = 0 nS, Vm_pre_ = −80 mV), the synaptic current reached −0.5 nA at a postsynaptic membrane potential of −65mV. In contrast, at a g_h_ of 60 nS, (Vm_pre_ = −71.5 mV) the same synaptic current was only achieved at −56.5 mV. This 8.5 mV shift in the postsynaptic membrane potential reflected the same 8.5 mV change of the presynaptic membrane potential that was caused by I_h_. Consistent with our previous data (Fig. 4C), we obtained no further changes to the behavior of the synaptic current when we additionally added I_h_ to the postsynaptic neuron so that it was present in both neurons (Fig. 4G, green).

In summary, the model predicts that by depolarizing the presynaptic membrane potential, I_h_ reduces the membrane potential difference between pre- and postsynaptic neurons. This reduces the synaptic current, and consequently the ePSP amplitudes. Only when the postsynaptic neuron depolarizes does the membrane potential difference increase, and concurrently do the synaptic current and ePSP amplitudes. Hence, the activation function of the synaptic current and the voltage dependence of the ePSPs shift to more depolarized postsynaptic membrane potentials.

### I_h_ depolarizes the MCN1 axon terminal and shifts the voltage dependence of the MCN1-LG postsynaptic currents

To test whether the model predictions held true in the biological system, we recorded simultaneously from the MCN1 axon terminal and LG. We first compared changes in axon membrane potential in saline (with I_h_) and in CsCl (no I_h_). We found that, as predicted, with I_h_ present, the membrane potential was more depolarized (Fig. 5A). We also noted that ePSP amplitudes changed in the predicted direction when the MCN1 axon membrane potential was manually altered. Fig. 5B shows an original recording of the MCN1 axon where a long hyperpolarizing current pulse was applied. During this pulse, the axon membrane potential slowly depolarized due to I_h_. Importantly, the hyperpolarization of the axon caused an immediate increase in ePSP amplitude in LG (outlined arrow in Fig. 5B), which was followed by a slow reduction in ePSP amplitude (red arrow) as the axon membrane potential continued to depolarize (see also inset of Fig. 5B). Finally, when we blocked I_h_ with CsCl (Fig. 5C) and repeated the same manipulation, the ePSP amplitude still increased when the axon was hyperpolarized. However, the slow depolarization of the axon membrane potential was gone, and so was the reduction in ePSP amplitude. These results are consistent with I_h_ causing a depolarization of the axon membrane potential, which in turn changes the voltage dependence of the ePSPs in LG.

**Figure 5.**
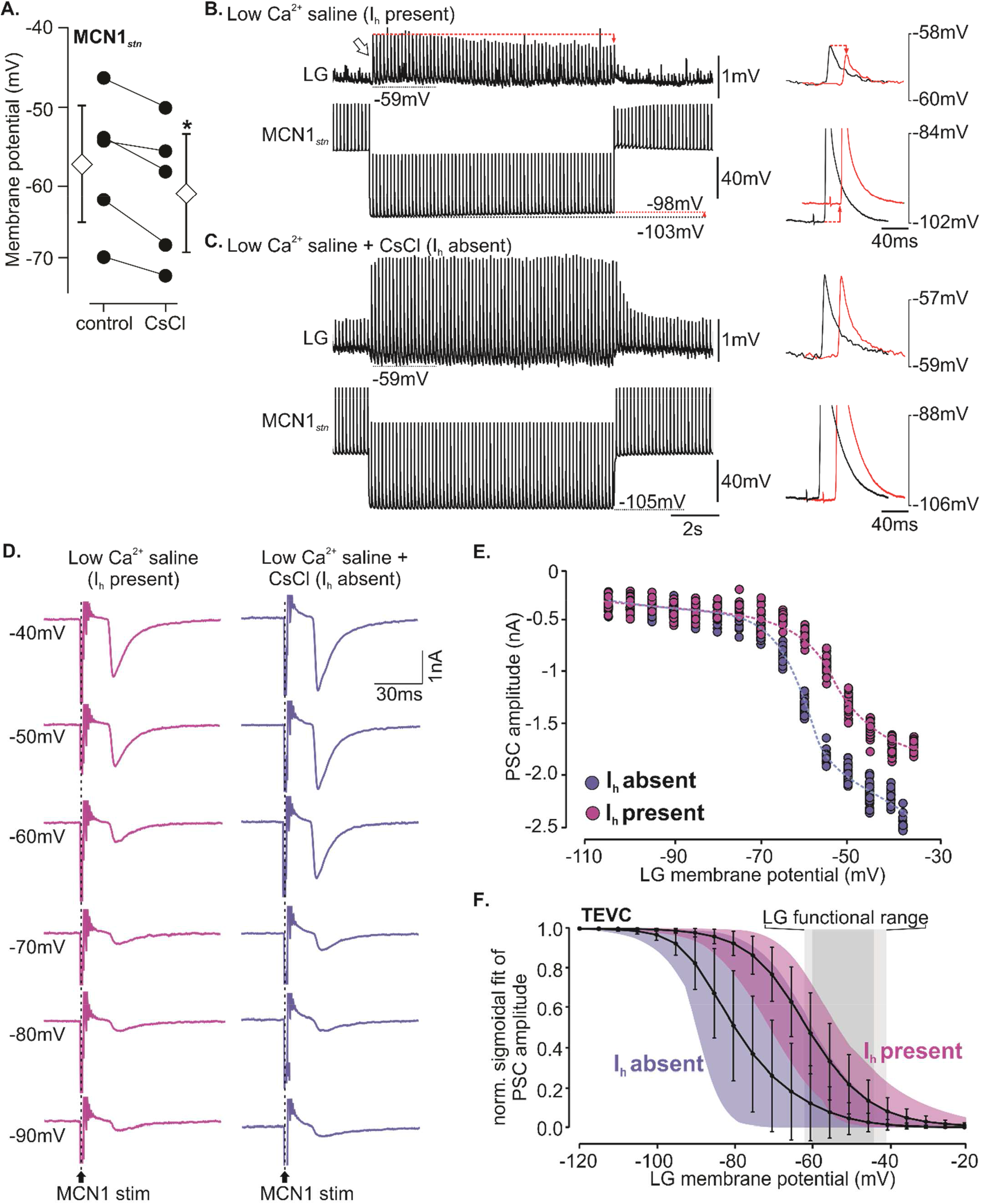
Presynaptic I_h_ shifts the voltage dependence of the MCN1 ePSPs toward the functional range of the LG membrane potential. A. The resting membrane potential of the MCN1 axon terminal hyperpolarized when I_h_ was blocked by CsCl. On average, the resting potential was −57.3 ± 8.1 mV in low calcium saline and −61.0 ± 8.2 with CsCl (N = 5; p<0.05, paired t-test). B. Simultaneous original recordings of LG and the MCN1 axon terminal. MCN1 was continuously stimulated at 5 Hz, and MCN1 was hyperpolarized for several seconds. Left: in low calcium saline, a sag potential slowly depolarized the trough membrane potential (red dotted line). Concurrently, the amplitude of the ePSPs in LG diminished (red dotted line and arrow). Outlined arrow: the hyperpolarization itself caused an increase in MCN ePSP amplitude. Right: magnification of MCN1 trough potential and action potentials (bottom traces), in addition to ePSPs in LG (top). One action potential and one ePSP from the beginning of the hyperpolarization are shown in black. One action potential and one ePSP from the end of the hyperpolarization are shown in red. In the latter, I_h_ had depolarized the trough membrane potential. C. Same manipulation as in B., but with I_h_ blocked by CsCl. The slow depolarization of the MCN1 membrane potential in response to hyperpolarization was absent. There was also no more decrease in ePSP amplitude during the MCN1 hyperpolarization. D. Original current traces of two-electrode voltage clamp LG recording, at different clamp potentials (−40 to −90 mV). Individual ePSCs were elicited through MCN1 axon stimulation with 1 Hz. Left: in low calcium saline. Right: with I_h_ blocked by CsCl. E. Quantification of synaptic currents obtained in the experiment shown in (A). Magenta: in low calcium saline. Purple: with I_h_ blocked by CsCl. F. Quantification of all experiments. The voltage dependence of the synaptic current of each experiment was fitted with a sigmoidal curve. Mean ± SD for all experiments are shown. Colored areas indicate the range of all observed data. Magenta: in low calcium saline. Purple: with I_h_ blocked by CsCl. The gray box indicates the functional range of LG’s membrane potential. Light gray indicates SD.

If I_h_ indeed acts presynaptically, i.e., if its effects on the LG ePSPs are mediated by its actions on the MCN1 axon terminal, we would expect to see a change of the synaptic current in LG. Indeed, the model (Fig. 4G) predicted that the observed activation of the synaptic current should shift towards more depolarized membrane potentials of the postsynaptic neuron (here: LG). We thus performed two-electrode voltage clamp on LG and measured synaptic currents during 1 Hz stimulation of the MCN1 axon in the *ion*. These experiments were carried out in low Ca^2+^ saline to suppress chemical synaptic transmission. We altered LG’s membrane potential between −20 and −120 mV. As predicted, the amplitudes of the electrical postsynaptic currents (ePSCs) behaved similarly to the ePSP amplitudes. Specifically, more depolarized membrane potentials caused larger ePSCs. The traces in Fig. 5D show original recordings of LG’s ePSCs at selected membrane potentials, with and without I_h_. Fig. 5E shows the quantitative analysis for this experiment. Like for the ePSPs, the increase in synaptic current with membrane potential was approximately sigmoidal in shape. Fig. 5F shows the averaged data from all animals tested (N=13). The average midpoint voltage was −60.4±6.0 mV and ePSCs increased notably in amplitude above −80 mV. They approached maximum values around −40 mV. Hence, some of the dynamic range of the synaptic current overlapped with LG’s functional range of membrane voltages (gray in Fig. 5F). When we blocked I_h_ with CsCl (Figs. 5D-F, purple), the midpoint of the ePSC amplitude change shifted to more hyperpolarized voltages, consistent with a presynaptic action of I_h_ on the voltage dependence of the electrical synapse and mirroring the model predictions (Fig. 4G). On average, the midpoint of the voltage dependence was now −77.6±9.6 mV (N=11, Mann-Whitney U-test, p<0.001, significantly different from no CsCl control). In summary, our data demonstrates that presynaptic I_h_ modulates the voltage dependence of the electrical synapse between MCN1 and LG by depolarizing the membrane potential of the MCN1 axon terminal.

## Discussion

Our data demonstrate that the modulation of the ePSP voltage dependence by I_h_ is a critical component of the MCN1-LG electrical synapse. It shifts the voltage dependence of the ePSPs to more depolarized voltages, allowing the dynamic range of the ePSP voltage dependence to overlap with the functional range of LG membrane potentials.

### I_h_ is critical to the function of the voltage dependence of the MCN1-LG ePSPs

In the MCN1 version of the gastric mill rhythm, the release of the neuropeptide CabTRP Ia from MCN1 is the main driver behind the slow bursting activity of LG [9, 18]. In contrast, the concurrent electrical PSPs that MCN1 elicits contribute only slightly to LG’s ability to burst. When chemical release from the MCN1 axon is blocked, and only ePSPs remain, LG can produce individual action potentials, but no regular bursting can be observed [9]. Instead, MCN1’s ePSPs contribute to the maintenance of the burst once it is created by prolonging burst duration and increasing spike number. Their voltage dependence is an important feature that supports this function. At resting membrane potential, as well as during the LG interburst, ePSP amplitudes are small. They increase slowly after burst start and reach their largest amplitudes towards the end of the burst. To achieve this, ePSP amplitude must increase in a range of membrane voltages that LG traverses between interburst and burst, and the increase must occur slowly so that only the later portions of the burst are affected. Our data shows that the time course of the ePSP amplitude change is indeed slow, on the order of several hundred ms (Supplemental Figure S1B (https://doi.org/10.6084/m9.figshare.17303918.v1)). We also demonstrate that I_h_ is a critical factor in allowing the ePSP voltage dependence to fulfill its function. With I_h_ blocked, ePSP amplitude was already close to maximum at LG’s resting membrane potential, and the dynamic range between resting membrane potential and spike threshold was small. In contrast, with I_h_ present, the dynamic range of the voltage dependence was more depolarized. This facilitated amplitude changes between resting membrane potential and spike threshold.

### Is the ePSP voltage dependence a gap junction property or caused by factors in the extra-junctional membrane?

When we first measured the time constant with which ePSP amplitudes changed, we were surprised by the extraordinarily slow change (several hundred ms, Supplemental Figure S1B (https://doi.org/10.6084/m9.figshare.17303918.v1)). This time constant is roughly on the time course of I_h_ in gastric mill neurons and initially suggested to us that I_h_ not just shifts the voltage dependence but is instead involved in creating it. Looking only at the typical voltage range in which LG operates seemed to further support this notion since ePSP amplitudes in this range are large and mostly unchanged when I_h_, is blocked. This suggested that blocking I_h_ also removed the voltage dependence. And indeed, there is evidence that voltage dependence of electrical synapses can arise from intrinsic membrane properties, such as spatially close voltage-gated ionic currents. Persistent Na^+^ conductances in particular have been implicated in endowing electrical transmission with voltage-dependent properties [4, 21–23]. Their slow inactivation kinetics make them ideal to alter electrical transmission postsynaptically, either through changing the extrajunctional membrane resistance or the membrane potential. In the auditory axon terminals of goldfish, for example, electrical PSPs are amplified by persistent sodium currents and diminished by potassium currents [22]. In cerebellar Golgi neurons, where the de- and hyperpolarizing components of action potentials reach neighboring cells through electrical synapses, the hyperpolarizing components dominate the electrical PSP when the receiving cell is at resting membrane potential. However, when depolarized, activation of the persistent sodium current amplifies the depolarizing PSP component, switching the PSP from predominantly hyperpolarizing to predominantly depolarizing [3, 4]. I_h_ has also been implicated in creating voltage dependence at an electrical synapse. Here, the effect was mediated by a decrease in membrane resistance in the extra-junctional membrane, which caused changes in the coupling coefficient of the gap junction [6]. In contrast to these examples, our data shows that the voltage dependence of the MCN1-LG ePSPs is still present and unchanged in its sigmoidal shape when I_h_ is blocked. The midpoint of the sigmoidal had shifted to more negative values. Hence, I_h_ could not have been pivotal to endow the electrical synapse with a voltage dependence. Our physiological data are also consistent with previous studies [9] and with our own model data that demonstrate that I_h_ in a presynaptic neuron can shift the midpoint of a voltage-dependent electrical synapse. Furthermore, when we voltage clamped LG, excluding extra-junctional influences in the postsynaptic cell, the synaptic currents showed a similar voltage dependence as the ePSPs, and their voltage dependence shifted to more negative voltages when I_h_ was blocked. Finally, the fact that imposed voltage changes in MCN1, as well as those elicited by I_h_, caused concurrent changes in ePSP amplitude further supports the conclusion that the observed voltage dependence is a property of the electrical synapse itself rather than being due to effects on the extra-junctional membrane. The model further supports this conclusion since changing the resistance of the presynaptic membrane had negligible effects on the voltage dependence.

### Modulation of ePSP voltage dependence by I_h_

Neurons possess a variety of heterogeneous ionic conductances, creating complex and highly non-linear intrinsic and synaptic dynamics, and there is mounting evidence that diverse sets of ion channels are distributed throughout all neuronal compartments, including axons and their terminals. Three main functions have been attributed to I_h_ [24]: (1) It contributes to the resting potential. Blocking I_h_ leads to a shift in the resting potential towards more hyperpolarized voltages. (2) It shapes neuronal responses to hyperpolarizing input. Due to its depolarizing effects, I_h_ can shape the amplitude and time course of PSPs, and thereby change the response of the cell to synaptic input. (3) It contributes to the pacemaker properties of a cell by causing a postinhibitory rebound that facilitates rhythmic activity.

Our data suggests that I_h_’s effect on the membrane potential is to cause a shift in the ePSP voltage dependence. Our MCN1 recordings show that changing the axon terminal membrane potential has a strong impact on the ePSP amplitude, and that such amplitude changes occur concurrently with membrane potential changes when I_h_ is activated by hyperpolarizing the terminal. Changes in ePSP amplitude also occur when I_h_ is blocked and the membrane potential is artificially altered, albeit without associated conductance change. Our model corroborates these conclusions and further suggests that changes in the strength of I_h_ can impact ePSP voltage dependence only when I_h_ is in the presynaptic neuron (MCN1). Here, changing the membrane potential was necessary and sufficient to elicit the shift of the voltage dependence.

Signal transmission at electrical synapses is complex and a host of neuronal and synaptic properties affect it, including differences in pre- and postsynaptic membrane resistances, capacitances, cell sizes, and membrane potentials [1]. The key to understanding how changes in membrane potential can affect the voltage dependence of the MCN1-LG ePSPs seems to be the rectification of this synapse. When both the presynaptic MCN1 axon terminal and the postsynaptic LG neuron are at rest, there is little voltage difference across the junction. Hence, junctional conductance is low and there is little driving force across that small junctional conductance. Incoming MCN1 action potentials thus only elicit small synaptic currents, yielding small ePSPs in the postsynaptic LG. When LG depolarizes and begins its burst, the voltage difference between LG and the MCN1 axon terminal increases and so does the junctional conductance. Incoming MCN1 action potentials then produce large voltage differences across the junction. These voltage differences occur rapidly and are too fast to alter the overall synaptic conductance (due to the slow kinetics, see Supplemental Figure S1B (https://doi.org/10.6084/m9.figshare.17303918.v1)), and hence cause the large ePSPs observed during the LG burst. I_h_ in the presynaptic MCN1 is a critical factor to maintain the appropriate voltage difference between the axon terminal and LG. Without I_h_, the MCN1 axon terminal hyperpolarizes, causing a voltage difference between the pre- and postsynaptic sides even when LG is not bursting. Consequently, junctional conductance is already high, and when action potentials arrive at the presynaptic terminal, they cause large synaptic currents that yiel large ePSPs with only small margins left for further amplitude increase when LG bursts.

### Where is I_h_ located?

We noted that while the model suggested that I_h_ must be presynaptic to exert its effects on ePSP voltage dependence, sag potentials were present in the MCN1 axon terminal and in LG. This suggests that I_h_ may be present on the pre- and postsynaptic side. However, we recognized that the voltages at which I_h_ was activated in LG were rather hyperpolarized and large current injections were required to reach these membrane potentials. During gastric mill rhythms, LG’s most hyperpolarized membrane potential was −59 mV, which is approximately 30 mV more depolarized than the voltages at which I_h_ was visibly activated. LG is the largest cell in the STG, with a cell body size of up to 100 μm, and if at all present, I_h_ is likely to be located in the LG neuropil (similar to other STG neurons). The hyperpolarized activation values could thus indicate a substantial space clamp issue in this large cell, where with increasing distance of the somatic electrode from I_h_’s location, current measurement degrades and, due to the electrotonic distance, more hyperpolarizing current at the recording site is necessary to cause corresponding voltage changes at the current source. This effect may be exaggerated by impedance changes within the LG neurites [25] that favor current flow from the neuropil toward the soma, but not vice versa. However, I_h_ has never been recognized in LG before, nor does it seem necessary to create rhythmic gastric mill activity [18]. It also has a rather low expression profile in LG [26]. One of the problems in studies with electrically coupled cells is that it is difficult to tell where observed effects take place. Depending on the coupling strength of the electrical synapse, the two cells may even act as a single physiological unit [27], at least at times. It is thus possible that I_h_ is exclusively located in the MCN1 axon terminals and our LG recording only picked up the reflection of the MCN1 I_h_. This notion is supported by the fact that LG’s postinhibitory rebound was typically not accompanied by action potentials. We observed postinhibitory rebound with spike generation predominantly in MCN1, suggesting that I_h_ was present in the presynaptic terminal. The sag potential was also more pronounced in MCN1 than in LG and was achieved with small current injections. Given the voltage dependence of the electrical synapse and its time constant, it seems unlikely that our current injection into the MCN1 axon hyperpolarized LG far enough to cause a rebound that would be large enough to backpropagate through the gap junction into the MCN1 axon terminal and to an area in MCN’s axon where spikes can be initiated. No previous data about I_h_ in MCN1’s axon terminal is available. However, we found a consistent, albeit small, hyperpolarization of the axon resting membrane potential when I_h_ was blocked, and the sag potential was absent. The small membrane potential change was likely due to the electrotonic distance between recording site and the axon terminal and gap junction. We recorded the MCN1 axon at the anterior entrance to the STG, and thus just outside of the STG neuropil. It is reasonable to assume that the gap junction to LG lies within the neuropil.

### Neuromodulation of the MCN1-LG electrical synapse

Membrane conductances like I_h_ are subject to intrinsic and extrinsic plasticity and have been shown to be the target of neuromodulatory transmitters. The STG is subject to extensive modulation by neuromodulators that reach the STG either in the form of neurohormones [28], or by being released from sensory cells and the axon terminals of the projection neurons [29]. Their presence alters circuit output or robustness [7, 30–33]. This opens the fascinating possibility that the voltage dependence of the MCN1-LG ePSPs could be rapidly altered by modulation of I_h_ in the MCN1 axon terminal or by activating other ionic conductances that alter the MCN1 terminal membrane potential. For example, I_h_ in the STG is modulated by serotonin, which depolarizes the activation curve of I_h_ by ~10 mV [34–36] and is released in the vicinity of the MCN1 axon terminals [37]. Similarly, Dopamine is present in the STG [38–42] and activating D_1_ receptors leads to a long-term increase in I_h_ [43, 44]. For example, Dopamine depolarizes the axon of the pyloric PD neuron through its excitatory actions on I_h_ [11, 43, 45]. While we did not examine this in our investigation, studying modulatory influences on the MCN1-LG electrical synapse is the next step in unravelling the complex dynamics between intrinsic cell properties, extrinsic influences and electrical synaptic transmission.

## Acknowledgments

We would like to thank Alexandra Yarger, Daniel Nuccio, Chris Goldsmith, Doug Schuweiler, Amanda Smith, Bangxia Suo and Dana Tilley for help on early modeling for this project, and their thoughts on voltage-dependent synapses. We would also like to thank Ulm University for their support of LYB’s research at Illinois State University.

Supported by NSF IOS 1755098, NSF IOS 1354932, NSF 1828136, DFG STE 937/8-1 and STE 937/9-1.

## Supplemental information

**Supplemental Figure S1.**
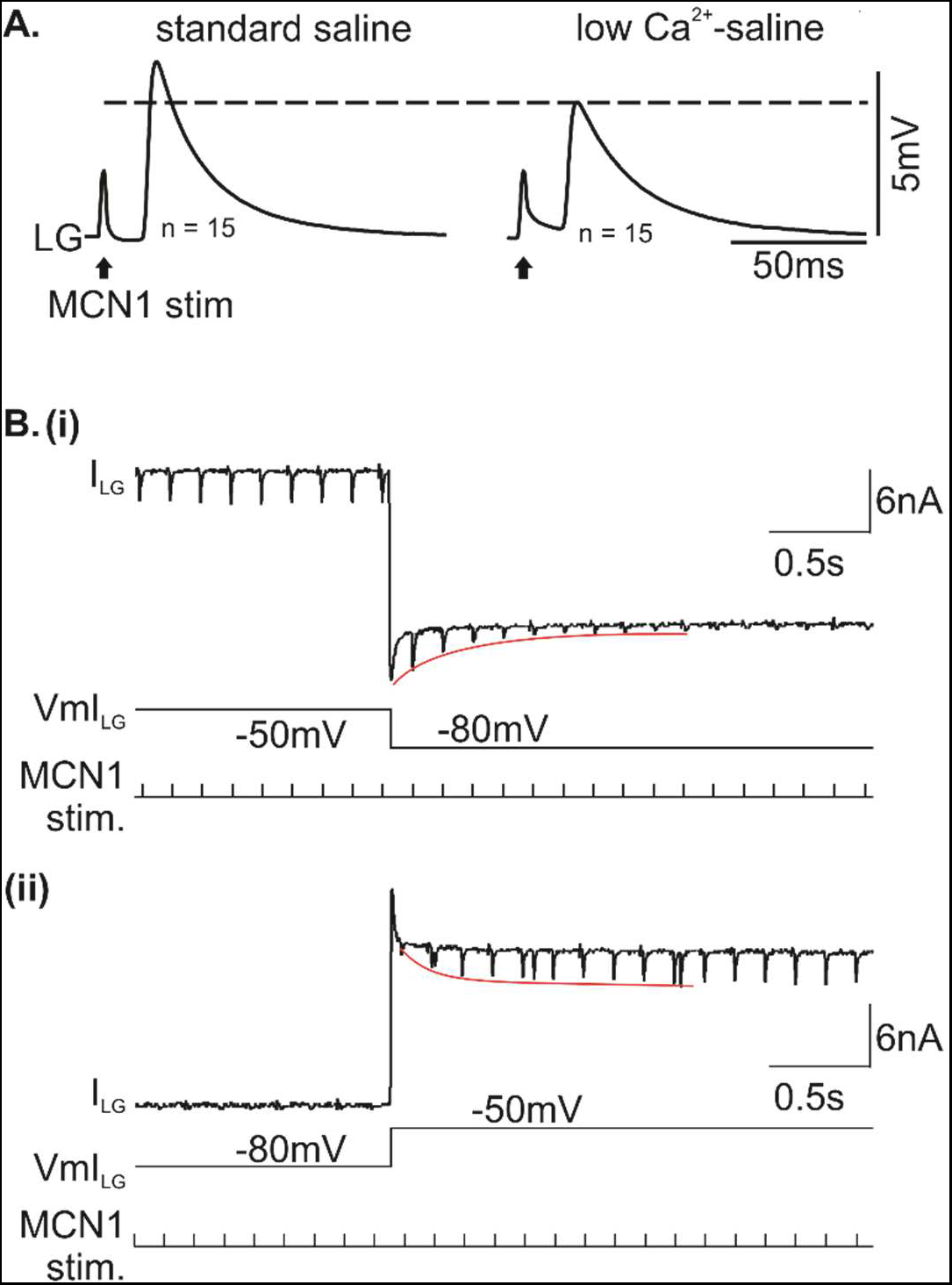
A. Time-triggered averages of MCN1 PSPs in LG. Individual ePSPs were elicited through MCN1 axon stimulation with 1 Hz. Left: in saline. Right: in low Calcium saline to block chemical synaptic transmission. While the amplitude diminished slightly, the major component of the PSP was electrical. PSPs were still present in low Calcium saline, demonstrating that he major component of the PSP was electrical. Nevertheless, PSP amplitude diminished, suggesting that a Calcium-mediated process enhances the PSP. B. Changes in ePSP amplitude occur with a slow time course. Two-electrode voltage clamp recording of MCN1 ePSCs in LG, in low Calcium saline. MCN1 was stimulated every 150 ms. (i) The LG membrane potential was hyperpolarized from −50 mV to −80 mV. (ii) The LG membrane potential was depolarized from −80 mV to −50 mV. Red lines show the decrement (i) and increase (ii) or the ePSC amplitudes after the voltage step. On average, the time constant for ePSC amplitude increase when LG was depolarized was 477 ± 281 ms (N=7). The average time constant for ePSC amplitude decrease when LG was hyperpolarized was 385 ±331 ms (N=6).

**Supplemental Table T1.**
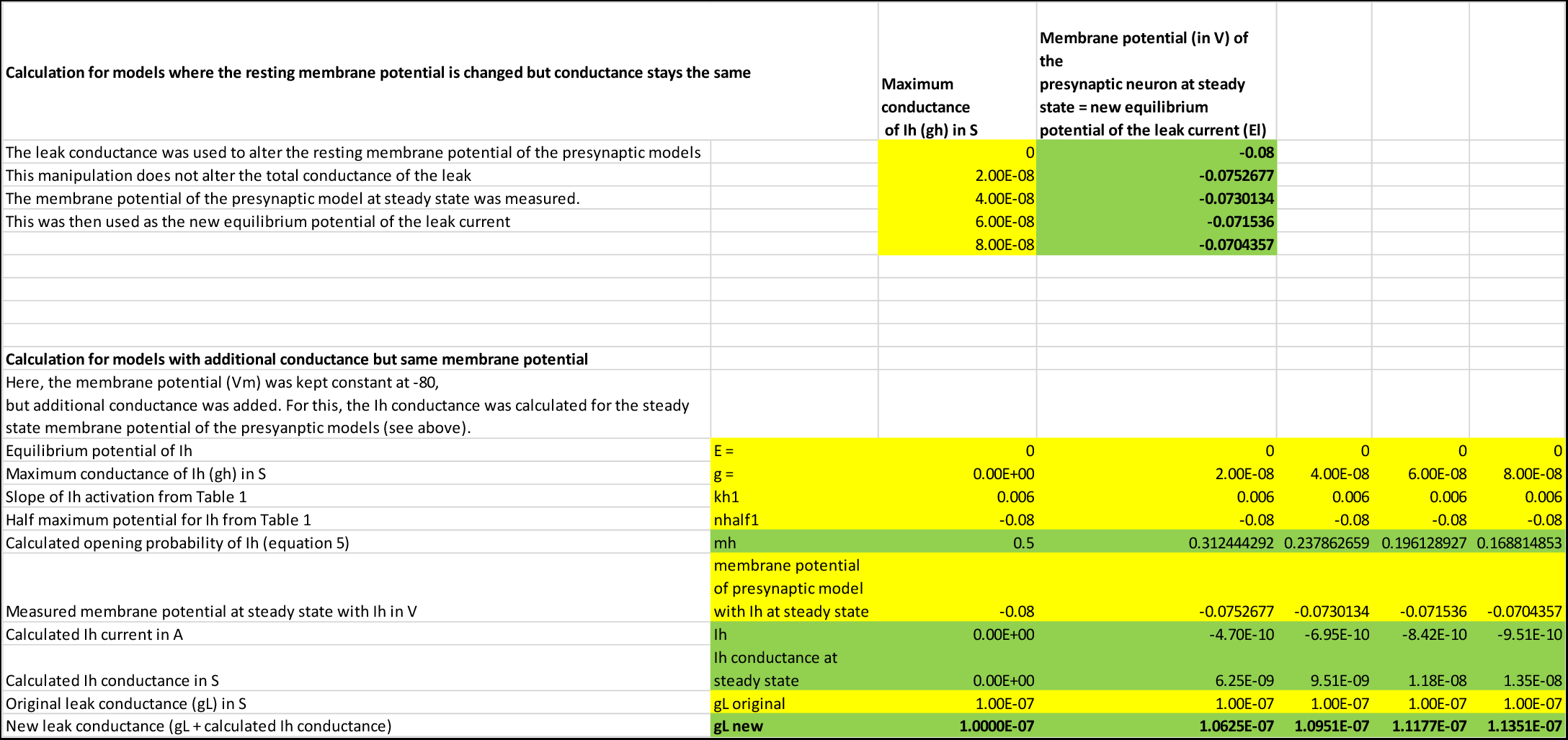
Top: measurement of membrane potential at steady state in models with different amounts of I_h_ (g_h_ was varied). Bottom: calculation of changes to leak conductance to mimic the effects of conductance increase by I_h_.

## Notes

### Competing Interest Statement

The authors have declared no competing interest.

### Summary of Updates

Revised to alter title and to clarify wording about the effects of Ih on gap junction signaling. A new panel was added to figure 4. Typos were corrected.

